# BRCA1 Hypermethylation In Sporadic Breast Cancers: Discovering A Novel Pathway To Tumorigenesis Via Coordinate NBR2 Deregulation And TNBC Transformation

**DOI:** 10.1101/2022.04.30.490082

**Authors:** Dipyaman Patra, Geetu Rose Varghese, Vishnu Sunil Jaikumar, Arathi Rajan, Neethu Krishnan, Krithiga Kuppuswamy, Rateeshkumar Thankappan, Priya Srinivas

## Abstract

Women with a family history of mutations in the Breast cancer susceptibility gene, BRCA1 will have an increased risk of developing breast neoplasms. However, majority of the breast cancers are sporadic where BRCA1 mutations are very rare. Instead, 5-65% of sporadic cases manifest BRCA1 promoter hypermethylation and 30-40% of such cases develop into Triple Negative Breast Cancers. Even then, the molecular mechanism of BRCA1 hypermethylation mediated breast tumorigenesis has remained an enigma till date. Here, we present a novel tumorigenesis pathway for breast cancers that engenders from BRCA1 hypermethylation by generating site-specific methylations in the BRCA1 promoter using a modified version of CRISPR technology.

We report that induction of site-specific methylation on BRCA1 promoter α effectuates a downregulation in BRCA1 expression via alteration in the balance between its alternate transcripts β and α. Induced BRCA1 hypermethylation is also responsible for the attenuation of a long noncoding RNA, NBR2 (Neighbour of BRCA1 gene 2), which is transcribed through the bidirectional BRCA1 promoter α in the reverse direction. Downregulation of NBR2 activates a feedback loop by leading to further downregulation of BRCA1 which is more evident under glucose starvation conditions and is associated with impaired DNA damage repair. BRCA1 hypermethylation also results in significant overexpression of β-hCG (human chorionic gonadotrophin), which was found to be associated with highly aggressive and drug-resistant forms of BRCA1 mutated breast cancers *invitro* & *in vivo* in our previous study. Further, we report a change in the hormone receptor levels as the tumor progresses which demonstrates how BRCA1 deficient cells modulate their expression of ER-α and ER-β to promote their proliferation in early stages of tumor development and at later stages, transform to a basal tumor subtype by shedding down the expression of ER-α & PR. Interestingly, we also discovered that modulation of ER-α expression upon BRCA1 hypermethylation is responsible for the alteration in BRCA1 transcript ratio. Finally, in *in vivo* mouse studies, BRCA1 hypermethylated tumors were found to be much larger, aggressive and invasive as compared to wildtype, BRCA1 and NBR2 knockdown tumors with downregulation of ER-α and PR; which explains the most probable reason behind high relapse rates in BRCA1 hypermethylated tumors.

**GRAPHICAL ABSTRACT:** 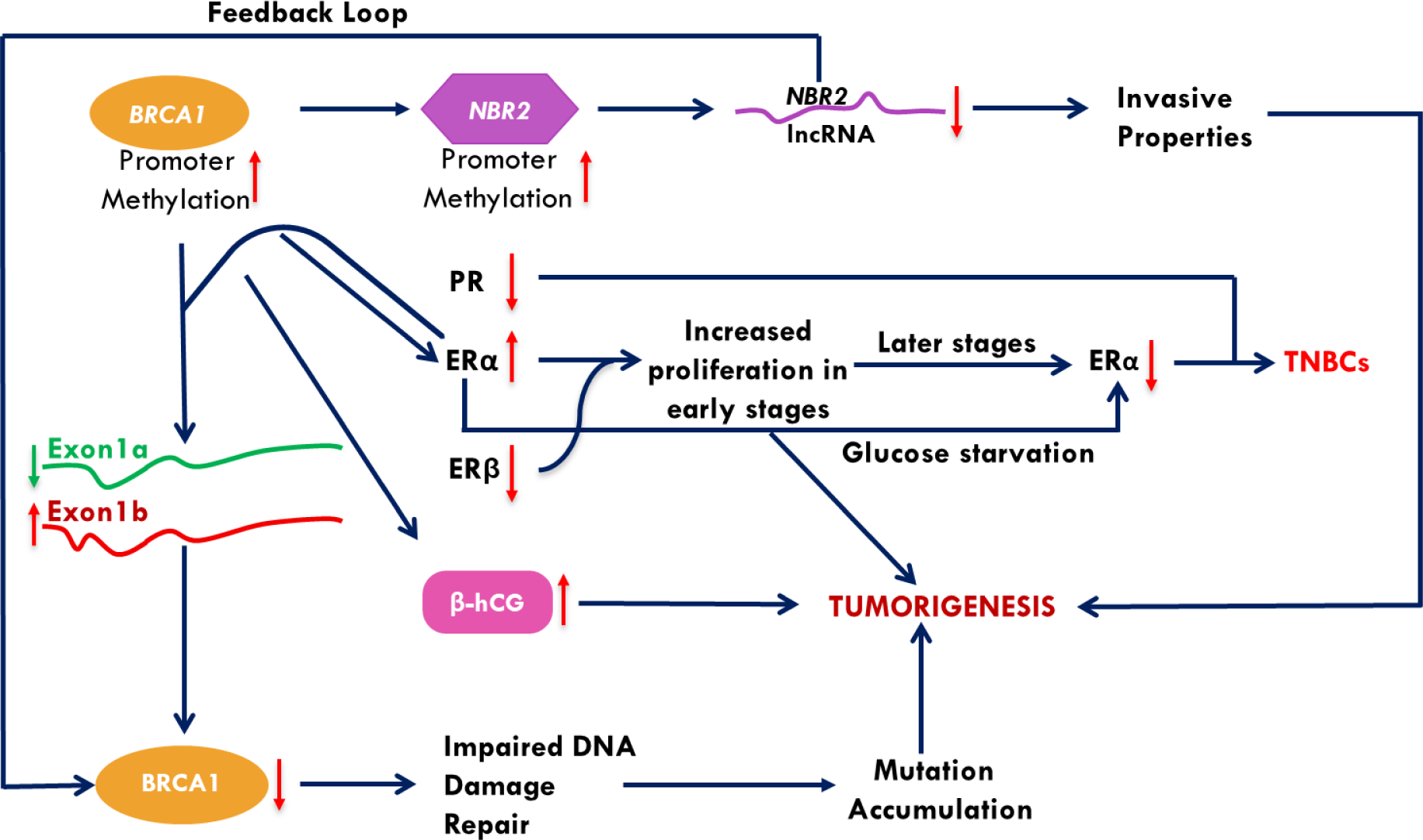

## INTRODUCTION

Breast cancer is the most common and dreadful form of cancer affecting females globally [1]. Alterations in the signalling of breast cancer susceptibility gene, BRCA1 is the impelling cause in many of these cases as it renders an important role in the accurate repair of DNA damage and maintenance of genomic integrity. Studies estimate that inherited mutations in BRCA1 can increase the cumulative risk of developing breast cancer with increasing age by 70-80% [2]. However, hereditary mutations account for only 5-10% of the total breast cancer burden while the rest of this menace can be attributed to sporadic cases. Interestingly, the story of BRCA1 mutations which is undisputed in inherited cases, don’t prevail when the cases are sporadic. This is attributed to the fact that most of the sporadic breast cancer patients manifest low BRCA1 expression without encountering a mutation in the gene [3, 4]. However, this loss of expression is reported to be the result of BRCA1 promoter hypermethylation which has been observed in sporadic breast cancer patients in a range from 5-65% [5–7]. Also, 30-60% of the Triple Negative Breast cancers (TNBCs) which respond very poorly to chemotherapy due to lack or low expression of Estrogen Receptor α (ER-α), Progesterone Receptor (PR) & Human Epidermal Growth Factor Receptor 2 (HER2) manifest BRCA1 hypermethylation associated with poor patient survival[8–13]. Several studies have also correlated BRCA1 promoter hypermethylation with low BRCA1 expression[11, 14]. But controversy still exists over the topic that whether loss of BRCA1 expression due to hypermethylation controls the process of tumorigenesis or is it a mere consequence of tumor progression.

The transcription of BRCA1 is controlled by two distinct promoters ‘α’ & ‘β’ leading to generation of two transcripts differing in their first exon (exon 1a in α and exon 1b in β transcript) [15]. The two promoters are activated at different levels in different tissues [15] which subsequently lead to differentially higher BRCA1 expression at sites of α-activation and vice-versa [16]. The presence of 3 non-canonical AUG codons upstream of the major translation initiation site in exon 1b at nucleotide positions 90, 109 & 134 (Fig 1) might be a major factor responsible for this differential expression as binding of the ribosome at the non-canonical start site in 5’ UTR can result in an alleviated engagement of translation machinery in the actual translation start site and a consequent lower translation efficiency of the β transcript [15]. Literature also shows significant downregulation of α and minor upregulation of β transcript in sporadic breast cancer [16]. All these facts indicate that the hypermethylation of BRCA1 promoter α could escort the process of tumorigenesis via deregulation of the principal α transcript and subsequent upregulation of β transcript.

**Fig 1:**
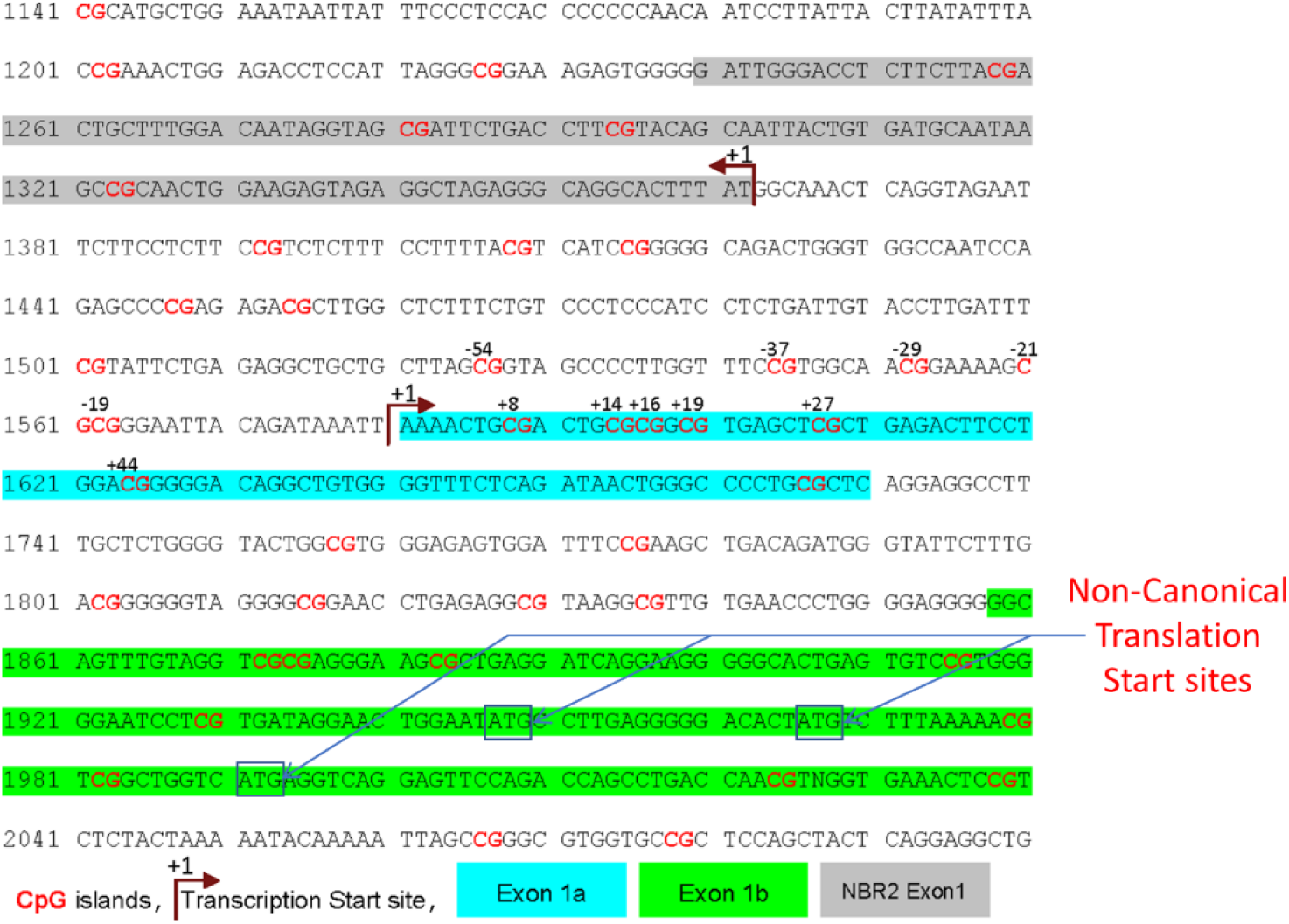
Position of BRCA1 Exon 1a (Blue) & 1b (Green), CpG islands (marked Red) and NBR2 Exon1 (Grey) [GenBank ID U37574]

Interestingly, BRCA1 promoter α is bidirectional in nature. Thus, it is responsible for transcription of another adjacent gene from the opposite DNA strand [17]. This gene, NBR2 (Neighbour of BRCA1 Gene 2), codes for a lncRNA which is mainly expressed under glucose stress conditions and regulates the activation of adenosine monophosphate-activated protein kinase (AMPK)[18]. By doing so, it represses tumor development via inhibiting anabolic processes like protein synthesis and promoting autophagy and cell survival under glucose stress conditions (Fig 2) [18]. A recent study has shown that anchorage-independent growth and *in vivo* xenograft tumor development is triggered in NBR2 knockdown condition [19]. Although literature survey reveals evidence for context dependent coordinate and reciprocal regulation of the two genes uncannily [20, 21], they have been found to be co-deleted in several breast and ovarian cancer patients [22, 23]. Also, one study demonstrated loss of promoter activity for both NBR2 and BRCA1 when the CCAAT box in BRCA1 promoter α was mutated [17]. Thus, it is implicative that the methylation of BRCA1 promoter α could potentially be a force driving the co-inhibition of BRCA1 and NBR2 in the process of breast tumorigenesis.

**Fig 2:**
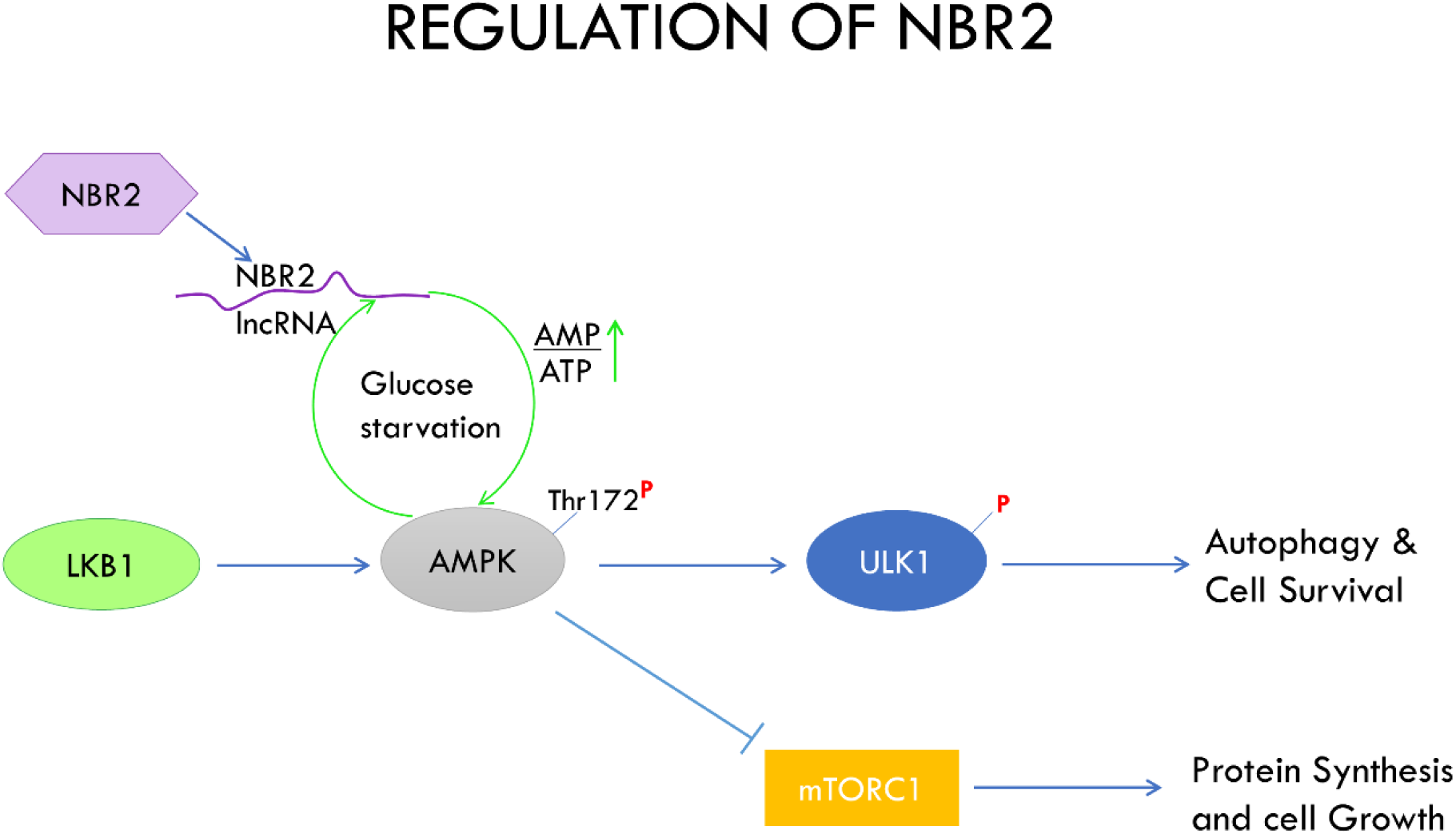
Regulation of NBR2 in AMPK pathway under glucose stress to supress tumor growth

**Fig 3:**
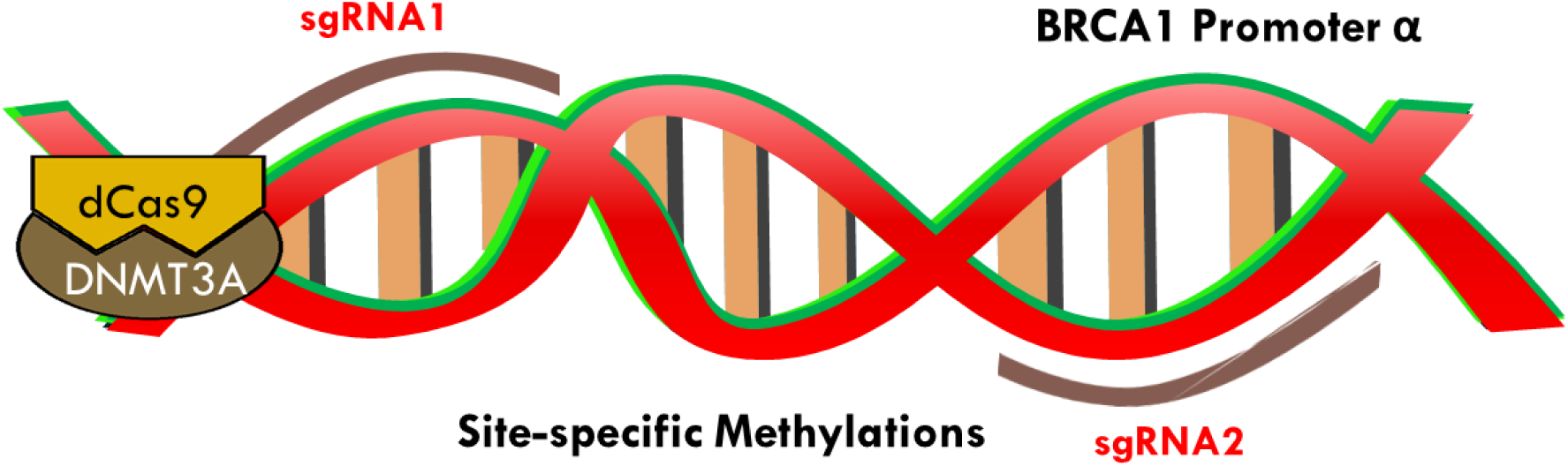
Induction of site-specific methylation on BRCA1 promoter CpG island of wildtype BRCA1 promoter α was targeted using a fusion construct of dead Cas9 and catalytic domain of DNMT 3A (*de novo* DNA methyl transferase) and two guide RNAs. The two guide RNAs span an area of <80bp and dictate the region where the fusion construct can bind to DNA and create site-specific methylations.

Another potential key player in the BRCA1 hypermethylation mediated breast tumorigenesis is the β subunit of Human chorionic gonadotropin (β-hCG). β-hCG has been implicated to have a pro-tumorigenic role in breast tumorigenesis. Our previous studies have demonstrated that wildtype BRCA1 can transcriptionally regulate β-hCG expression and knockdown of BRCA1 can lead to its overexpression. Moreover, the ectopic expression of β-hCG in BRCA1 mutated breast cancers results in aggressive breast tumours and can also lead to the induction of drug resistance [24, 25]. Interestingly, one of our recent studies also found β-hCG overexpression to be associated with BRCA1 promoter hypermethylation in Gestational Trophoblastic Diseases (GTD) and reversal of this hypermethylation via plumbagin led to the re-expression of BRCA1 and repression of β-hCG [26].

All this evidence indicates that BRCA1 promoter hypermethylation may have a pertinent role in sporadic breast tumorigenesis. Hence, we studied the role of BRCA1 hypermethylation in breast tumorigenesis by inducing specific methylations at the CpG island of wildtype BRCA1 promoter α using a modified version of CRISPR technology (Fig3) and demonstrated that hypermethylation of BRCA1 leads to low expression of BRCA1 via modulation of its alternate transcripts. BRCA1 Hypermethylation is responsible for augmentation of tumorigenesis process by deregulation of NBR2 and β-hCG. We also discovered that BRCA1 promoter hypermethylation causes the transformation of luminal type MCF7 cells to a basal subtype.

## METHODOLOGY

### Cell lines

The breast cancer cell lines MCF-7, MDA-MB-231, and viral packaging cell line HEK 293T used for the present study were collected from the RGCB Central Cell Repository facility. The cells were cultured in DMEM medium supplemented with 10 % FBS (GIBCO) and 100IU/ml Pen-Strep cocktail (Invitrogen). The cells were maintained in a humidified incubator with 5% CO2 at 37°C.

### Cloning of shRNAs/sgRNAs

For generating knockdown cell lines for BRCA1, NBR2, and ER-α, shRNA sequences were designed against the target mRNA sequences taken from Ensembl (https://www.ensembl.org) using the Kay Lab siRNA/shRNA/Oligo Optimal Design tool (siRNA/shRNA/Oligo Optimal Design stanford.edu).

For generating BRCA1 methylated cell lines target CpG island sequences were obtained from NCBI (GenBank ID U37574) and entered into the MIT sgRNA design software (http://crispr.mit.edu/) and sgRNA were selected based on location relative to other sgRNAs. The shRNA and sgRNA oligos were annealed and cloned into the vector, pLVTHM which was a gift from Didier Trono (Addgene plasmid #12247; http://n2t.net/addgene:12247; RRID: Addgene_12247). Details of shRNA and sgRNA sequences are in Table S1.

### Generation of Lentiviral particle

Lentiviral particles were generated in HEK 293T cell lines with lentiviral plasmids psPax2 and pMD2.G which was gifted from Didlie Trono (Addgene plasmid # 12260; http://n2t.net/addgene:12260; RRID:Addgene_12260 & Addgene plasmid # 12259; http://n2t.net/addgene:12259; RRID:Addgene_12259) using Lipofectamine TM 3000 (Catalog No. L3000008). The virus containing supernatants were collected at 24 hours and 48 hours post transfection and centrifuged at 3000rpm for 10 mins at 4°C to pellet down any residual cells and debris. The viral supernatant was then aliquoted and stored at −80.

### Generation of stable BRCA1 hypermethylated cell lines

Breast cancer cell line MDA-MB-231 and MCF 7 were grown and seeded onto a 60mm dish to be 70% confluent. After 24 hours the lentiviral particles were transduced to the target cell lines with 8μg/ml polybrene (Sigma TR-1003-G). For generation of stable methylated cells under the control of a tet-off promoter, target cells were co-transduced with lentiviral particles of TetO-dCas9-D3A [a gift from Grant Challen (Addgene plasmid # 78254; http://n2t.net/addgene:78254; RRID:Addgene_78254) and pCL-CTIG [a gift from Inder Verma (Addgene plasmid # 149021; http://n2t.net/addgene:149021; RRID:Addgene_149021)]. The eGFP and mCherry positive population were sorted using two filters by flow cytometry using BD FACSAria III Cell Sorter (BD Biosciences). The cells were then infected with lentiviral particles of either sgRNA1 or 2 or both and treated with 1μg/ml of doxycycline for 12 hours to stop the eGFP signal from pCL-CTIG. The eGFP positive population representing sgRNA positive cells were finally sorted by flow cytometry to establish the stable transduced cell lines.

### Methylation Specific qPCR Analysis (MS-qPCR)

Genomic DNA was isolated using QiAamp DNA Mini Kit (Qiagen, 51304) from cell lines following manufacturer’s protocol. Bisulfite conversion was done using Premium Bisulfite Kit (Diagenode, C02030030). Methylation analysis was done by qPCR using Methylation-specific PCR-Episcope MSP Kit (Takara Bio, R100B). EpiTect PCR Control DNA Set (Qiagen, 59695) was used to generate methylation standard curve and calculate the percentage of methylation in target cell lines. The details of primers used are given in Table S2.

### Real Time PCR (qRT-PCR)

mRNA was isolated using RNA isolation kit (Roche,11828665001) from cell lines. cDNA was prepared using High-Capacity cDNA Reverse Transcription (Life Technologies). qRT-PCR was performed using the SensiFAST™ SYBR® Lo-ROX Kit (Bioline, BIO-94005) on a QuantStudio^TM^ 5 Real-Time PCR system (Applied Biosystems^TM^, A28574). Gene expression data was normalized to the housekeeping gene GAPDH. The details of primers used are given in Table S2.

### Western Blot

To analyse the protein expression, immunoblotting was carried out for different proteins in the whole cell lysates of cell lines as described earlier [27]. The details of Antibodies used in this study are given in Table S3.

### TCGA data analysis

cBioportal for cancer genomics was utilized to analyse and correlate the expression and methylation levels of BRCA1, NBR2, ESR1, PGR and ERBB2 in breast cancer tissues samples (http://www.cbioportal.org/)

### Statistical Analysis

At least four *in vitro* experiments and four in vivo experiments were independently carried out and the data expressed as mean±SD. The probability of significant differences between two or more groups were analyzed by non – parametric Student’s t test or ANOVA. The GraphPad Prism (Version 8.0) software was used for all statistical analysis; appropriate tests are indicated in the figure legends. Statistical significance was defined as (*) p ≤ 0.05, (**) p ≤ 0.01, (***) p ≤ 0.001 and (****) p ≤ 0.0001.

## RESULTS

### Methylation of BRCA1 promoter modulated the balance between its alternative transcripts and effectuated BRCA1 downregulation

11 CpG at BRCA1 promoter α sites are known to be hypermethylated in sporadic breast tumors with low expression of BRCA1 protein [11, 12, 14] (Fig 1) and was the target region for site specific-specific methylation in our study. We used two target sgRNAs and a fusion construct of deadCas9 and DNMT3A to generate stable methylated cell lines under the control of a tet-off promoter (Fig 4). We confirmed the induction of methylation by MS-qPCR and found the maximum methylation in cells transduced with both sgRNA 1&2 (Fig 5).

**Fig 4(I):**
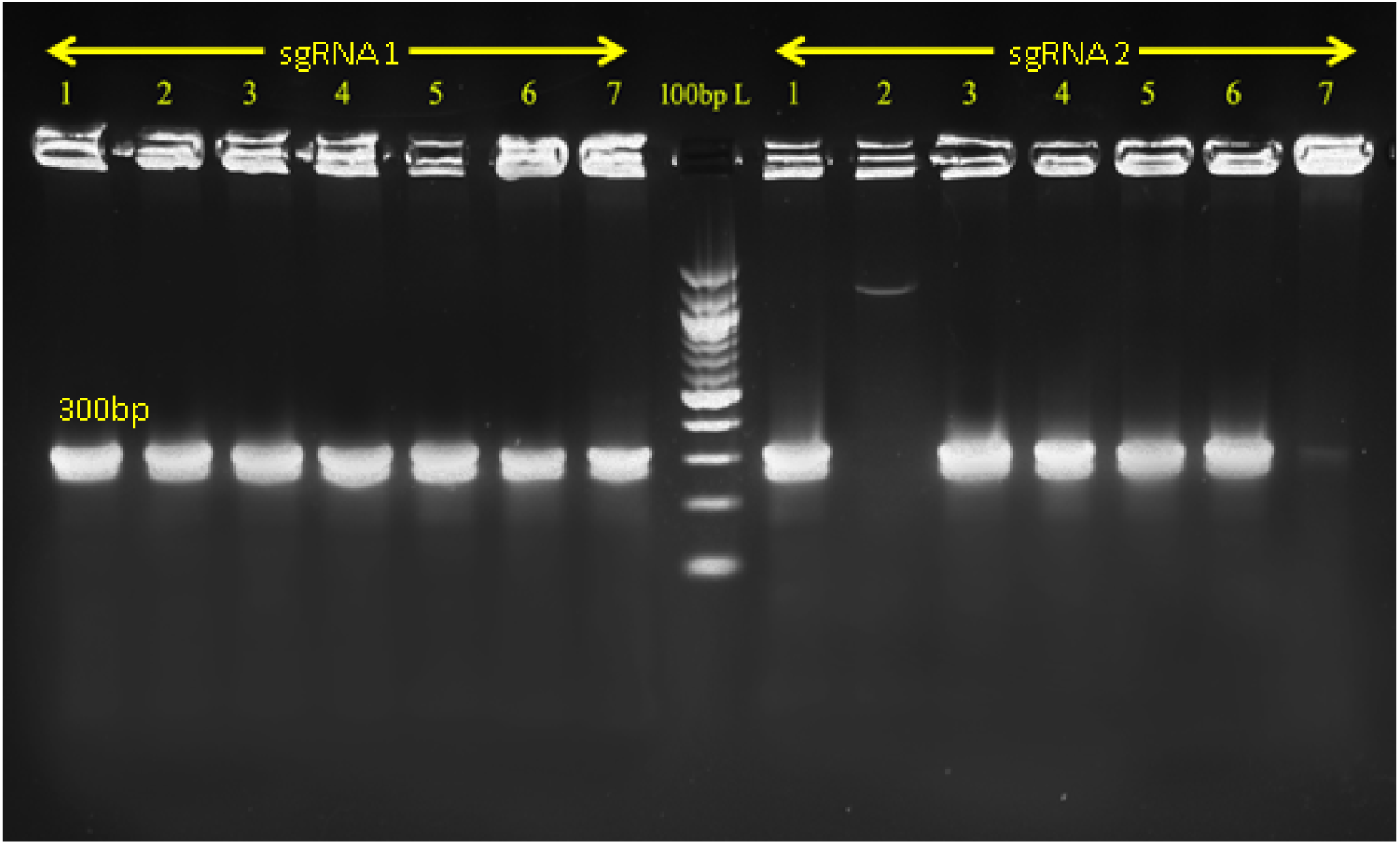
Colony PCR of two different sgRNA clones targeting the CpG island of BRCA1 Promoter α transformed into *stbl3* competent cells

**Fig 4(II):**
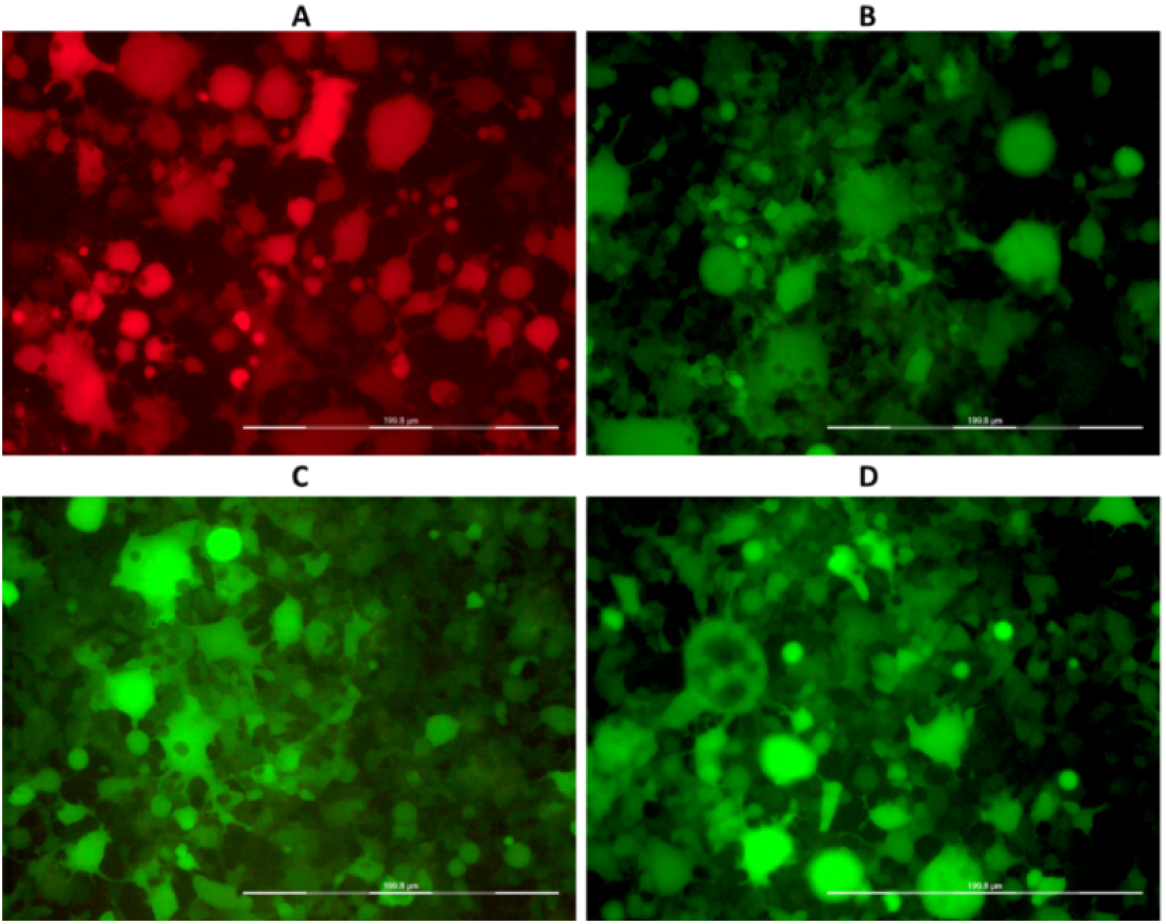
Lentiviral transfection of A) TetO-dCas9-Dnmt3a B) Empty sgRNA vector C) sgRNA1 D) sgRNA2 in HEK293T cells

**Fig 4 (III):**
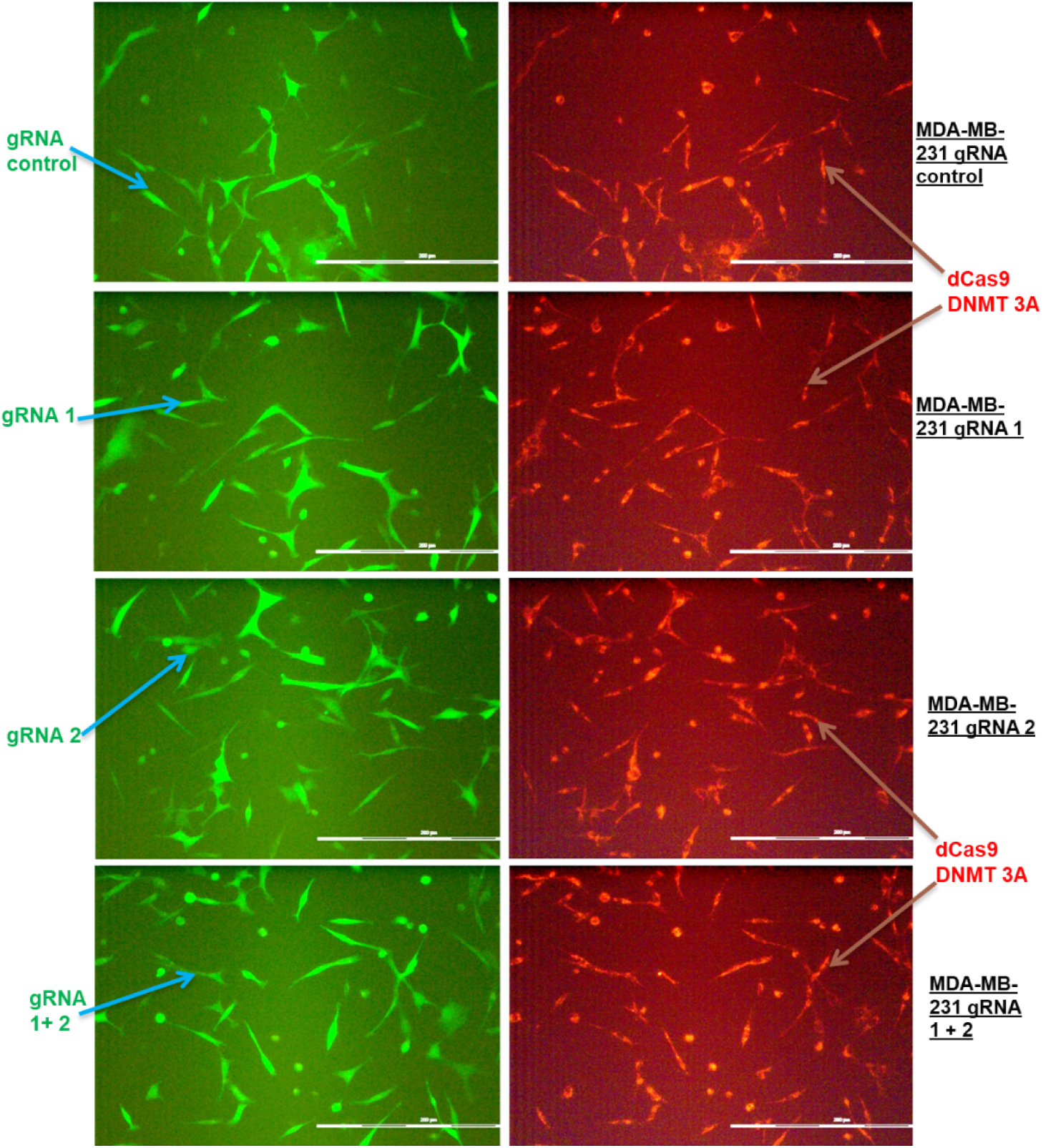
Generation of stable cell line after FACS sorting following transduction of two different lentivirus containing TetO-dCas9-Dnmt3a and sgRNA clones in MD-AMB 231 cells.

**Fig 5:**
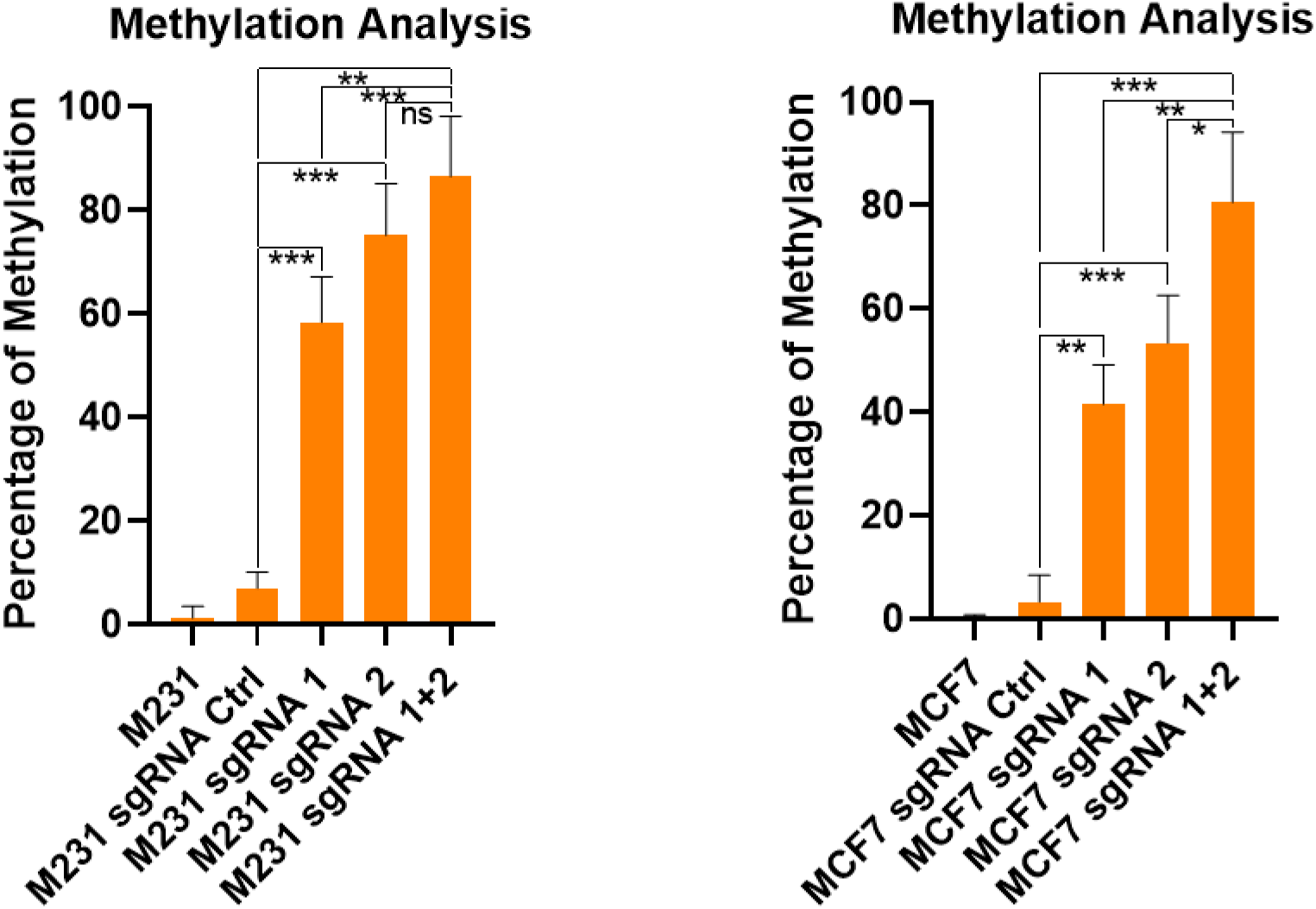
Methylation levels in Stable transduced cell lines analyzed by Methylation Specific qPCR in MDA MB 231 (M231) and MCF7 cells.

Our preliminary analysis of breast cancer datasets of TCGA showed an alleviation in BRCA1 expression with increasing levels of BRCA1 methylation in breast cancer patients (Fig 6). To confirm that our BRCA1 hypermethylation induced cells have the same effect on BRCA1 expression, we analyzed BRCA1 expression levels in the stable transduced cell lines. We found BRCA1 to be downregulated at RNA (Fig 7a) as well as protein level (Fig 7b) which was most significant in the cells with both sgRNAs and was more pronounced than BRCA1 knockdown (shBRCA1 cells) as well.

**Fig 6:**
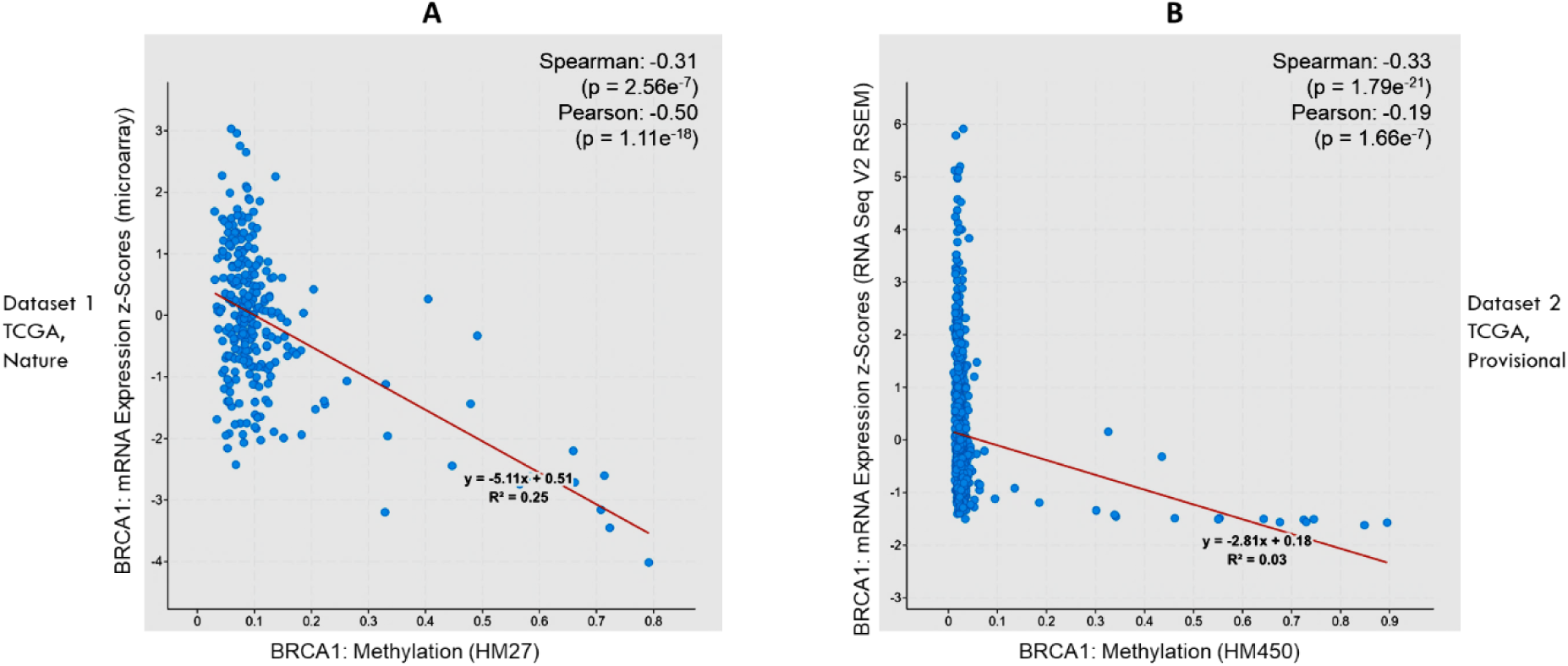
Plot A & B represent BRCA1 methylation vs BRCA1 mRNA levels from datasets Breast Invasive Carcinoma (TCGA, Nature 2012) and Breast Invasive Carcinoma (TCGA, Provisional) respectively.

**Fig 7:**
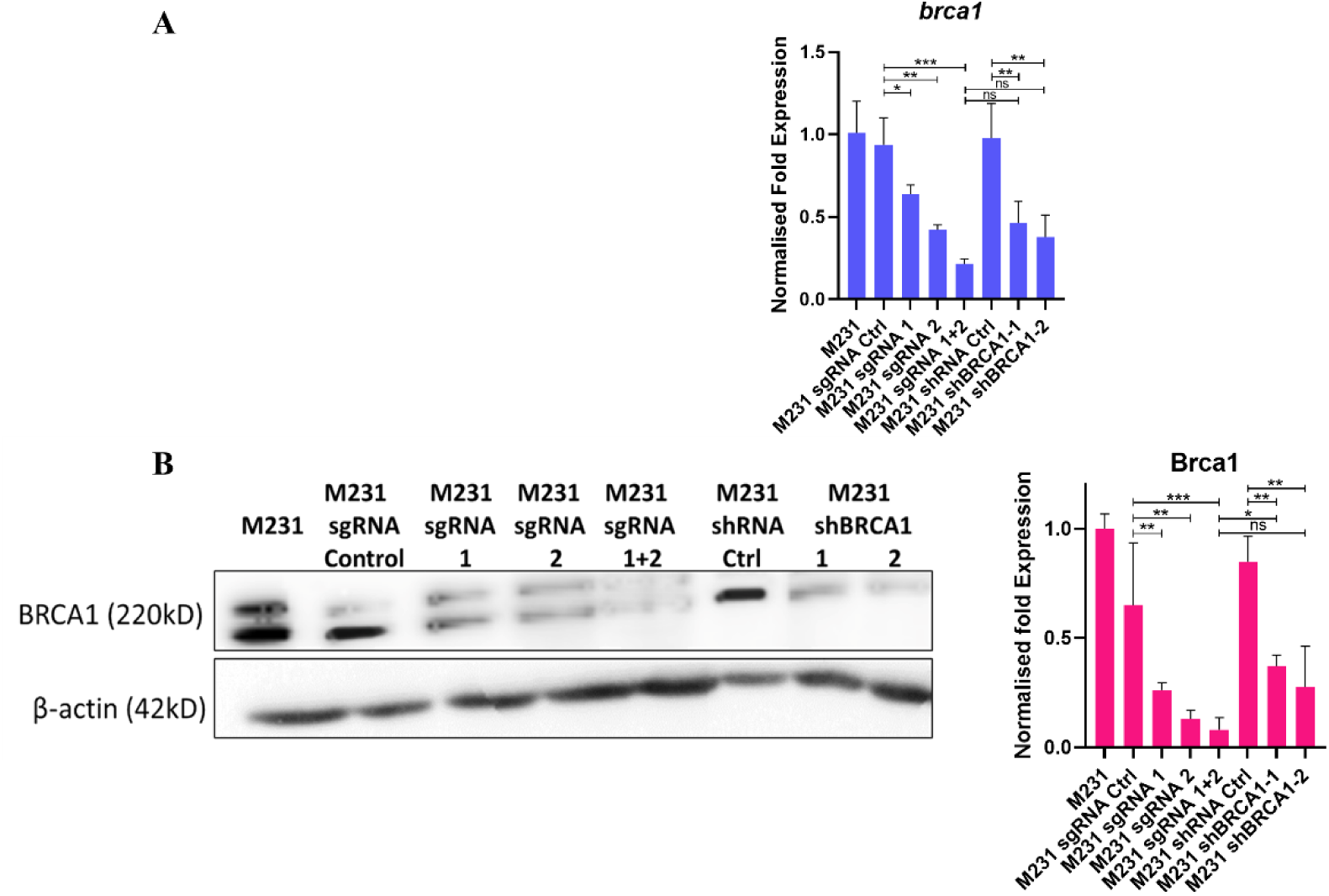
BRCA1 expression analysis at **(A)** mRNA (qRT PCR) & **(B)** protein level (western blot) in M231 cells show a downregulation of BRCA1 expression in BRCA1 hypermethylated condition.

Having confirmed BRCA1 downregulation upon BRCA1 promoter hypermethylation, we checked for the individual BRCA1 transcripts ‘α’ and ‘β’in these cell models. We found low levels of α and β transcript in both BRCA1 methylated and BRCA1 knockdown cells (Fig 8A). However, the β transcript levels were higher in the methylated cells when compared with the BRCA1 knockdown condition (Fig 8B). As a result, the ratio of transcript β to α was higher in methylated cells as compared to the control which was most pronounced in the cells with both sgRNAs 1&2 (Fig 8C). Simultaneously, this ratio was found to decline in BRCA1 knockdown cells (Fig 8C); thereby suggesting that the alternative BRCA1 transcripts get modulated differently under different conditions even when the total BRCA1 mRNA level is reducing. Altogether, we concluded that the induction of hypermethylation in BRCA1 promoter α effectuated a downregulation in its expression by altering the ratio of its alternate transcripts β and α.

**Fig 8:**
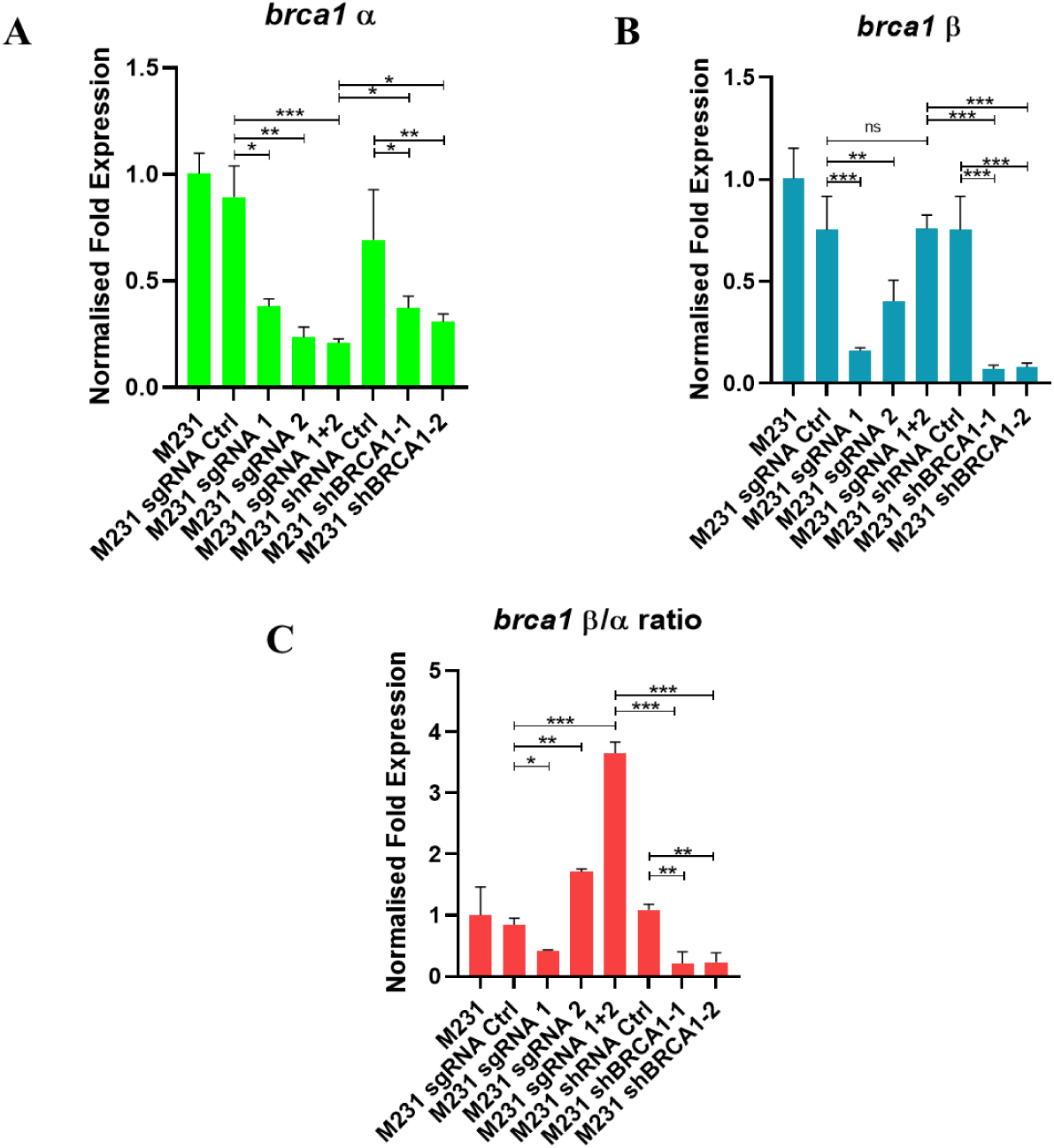
qRT-PCR analysis of individual BRCA1 transcripts **A)** α **B)** β and **C)** relative β/α transcript ratio reveals a modulation in alternate transcript level in BRCA1 promoter α hypermethylated MDA-MB-231 cells lines

### BRCA1 hypermethylation leads to the downregulation of lncRNA NBR2

Since NBR2 and BRCA1 share the same promoter (BRCA1 promoter α) we analyzed how the hypermethylation of BRCA1 promoter affected the NBR2 expression. We found diminished levels of NBR2 in methylation induced cells which was most significant in cells transduced with both sgRNA 1&2 (Fig 9A). However, the NBR2 expression remained unchanged in BRCA1 knockdown cells (Fig 9A), which insinuates that although BRCA1 doesn’t regulate NBR2 expression, methylation of BRCA1 promoter affects NBR2 transcription directly.

**Fig 9:**
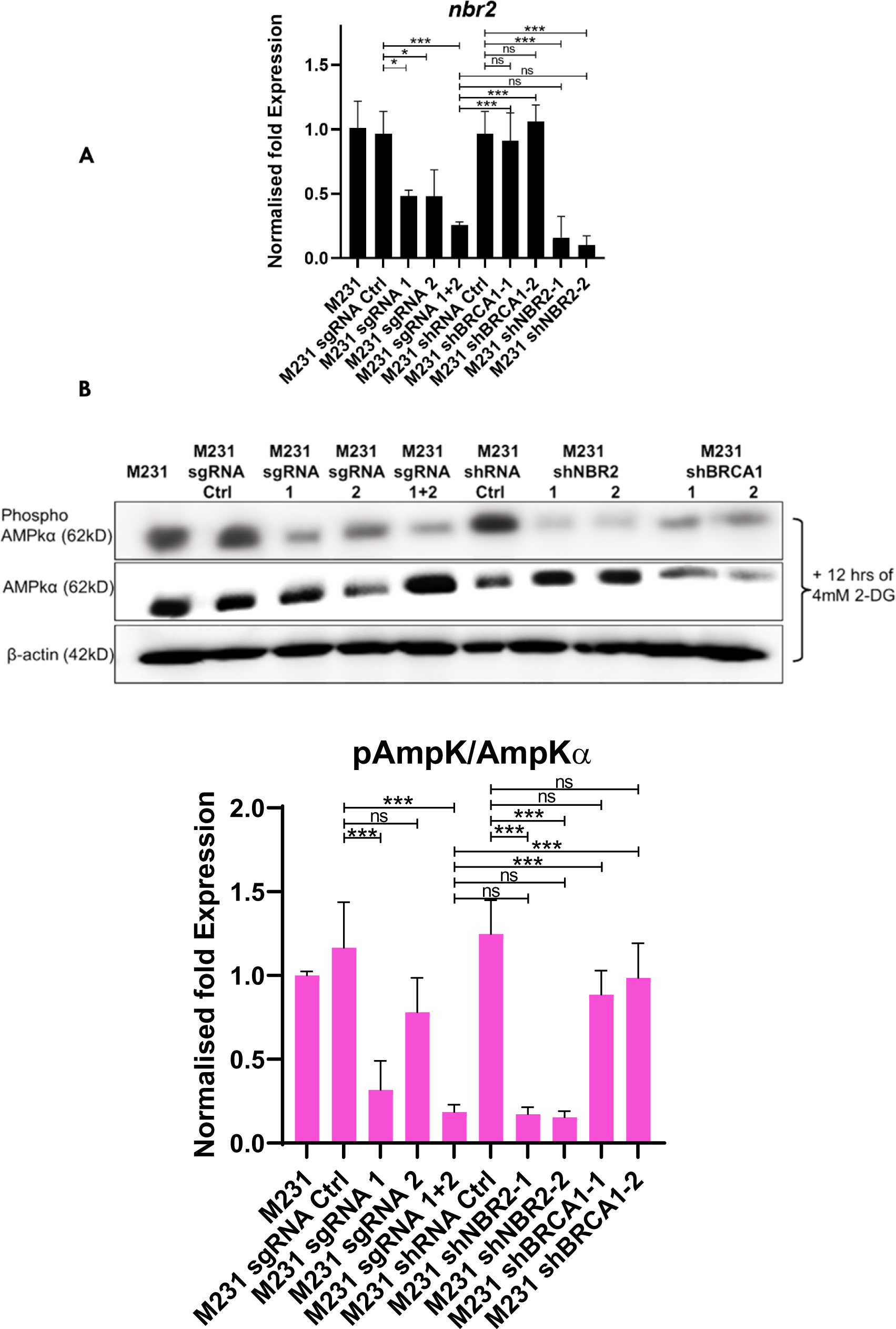
**(A)** Investigation of M231 cells show downregulation of lncRNA NBR2 at RNA level and **(B)** reduced phosphorylation of AMPKα (direct downstream target of NBR2) in BRCA1 promoter α hypermethylated condition which is absent in BRCA1 knockdown condition M231 cells.

Since NBR2 does not code for any protein, a direct confirmation of this result at protein level was not possible. As an alternative, we checked for AMPK phosphorylation which gets directly affected by NBR2 expression [18]. We performed western blot for phospho-AMPK and total AMPKα with NBR2 knockdown cells as control. Similar to qRT-PCR results, we found reduced levels of AMPK phosphorylation in BRCA1 methylated as well as NBR2 knockdown cells which was not observed in BRCA1 knockdown cells (Fig 9B). We also confirmed this result in two breast cancer datasets of TCGA via cBioportal which showed a positive correlation between NBR2 and BRCA1 hypermethylation (Fig 10 A & B) and a strong negative correlation between BRCA1 hypermethylation and NBR2 expression (Fig 10 C & D).

**Fig 10:**
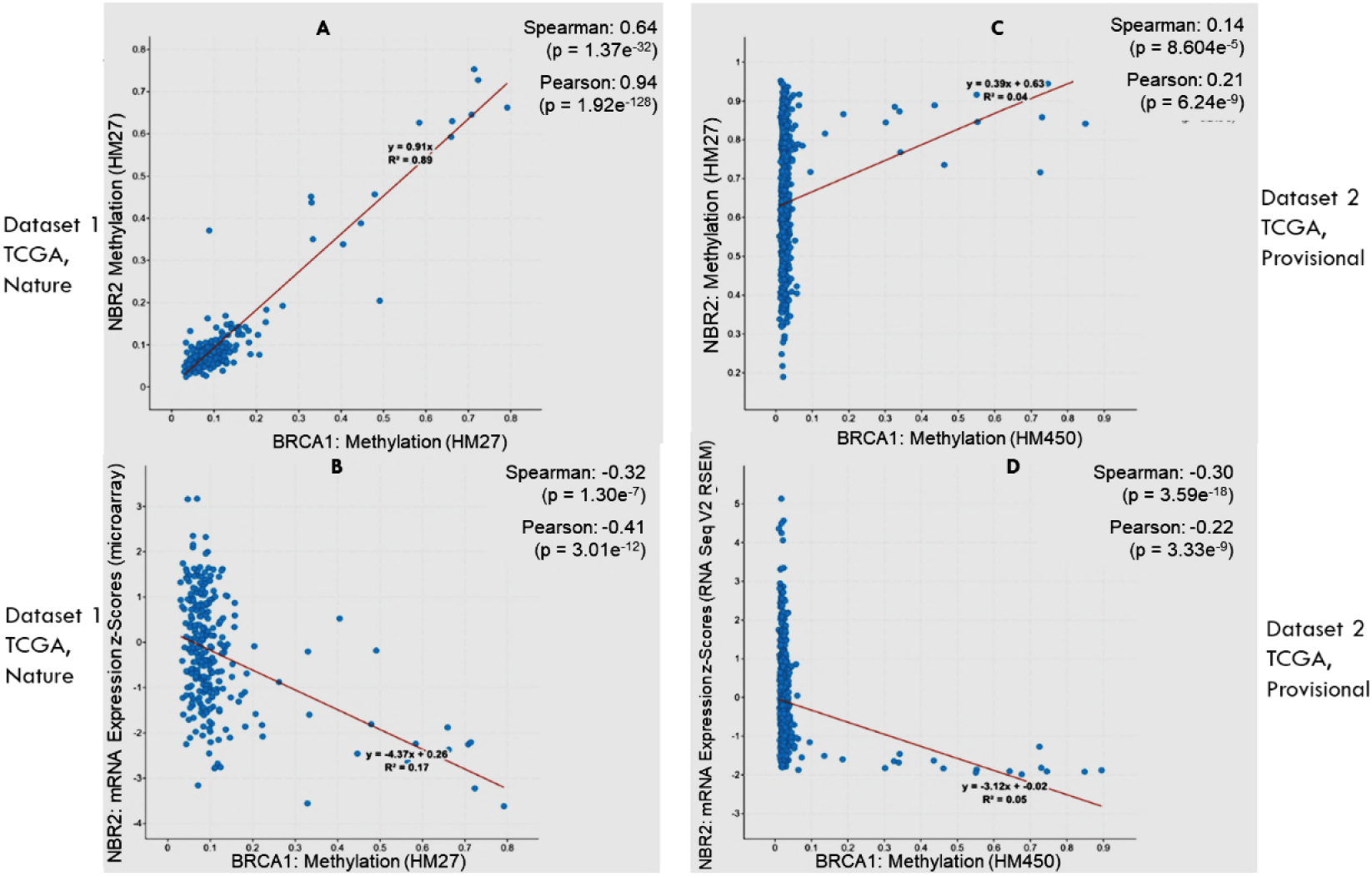
Plot A & B represent BRCA1 methylation vs NBR2 methylation from datasets Breast Invasive Carcinoma (TCGA, Nature 2012) and Breast Invasive Carcinoma (TCGA, Provisional) respectively. Plot C &D represent BRCA1 methylation vs NBR2 expression from the same datasets as A &B.

### NBR2 downregulation activates a feedback loop leading to further downregulation of BRCA1

Since, we observed NBR2 downregulation as an effect specific to BRCA1 promoter hypermethylation and not BRCA1 knockdown we did comparative analysis with NBR2 knockdown as control. Interestingly, we found diminished levels of BRCA1 in NBR2 knockdown cells which was more significant in double knockdown of BRCA1 and NBR2 (Fig 11 A). Western blot analysis showed that this downregulation was further pronounced under glucose starvation condition generated by treatment with 4mM 2-DG for 12hrs (Fig 11 B), as NBR2 is known to be transcribed prominently under glucose stress condition. Hence, we concluded that NBR2 downregulation engendering from promoter α methylation leads to the activation of a feedback loop resulting in further reduction of BRCA1 levels and augmenting the tumorigenic effect of BRCA1 hypermethylation.

**Fig 11:**
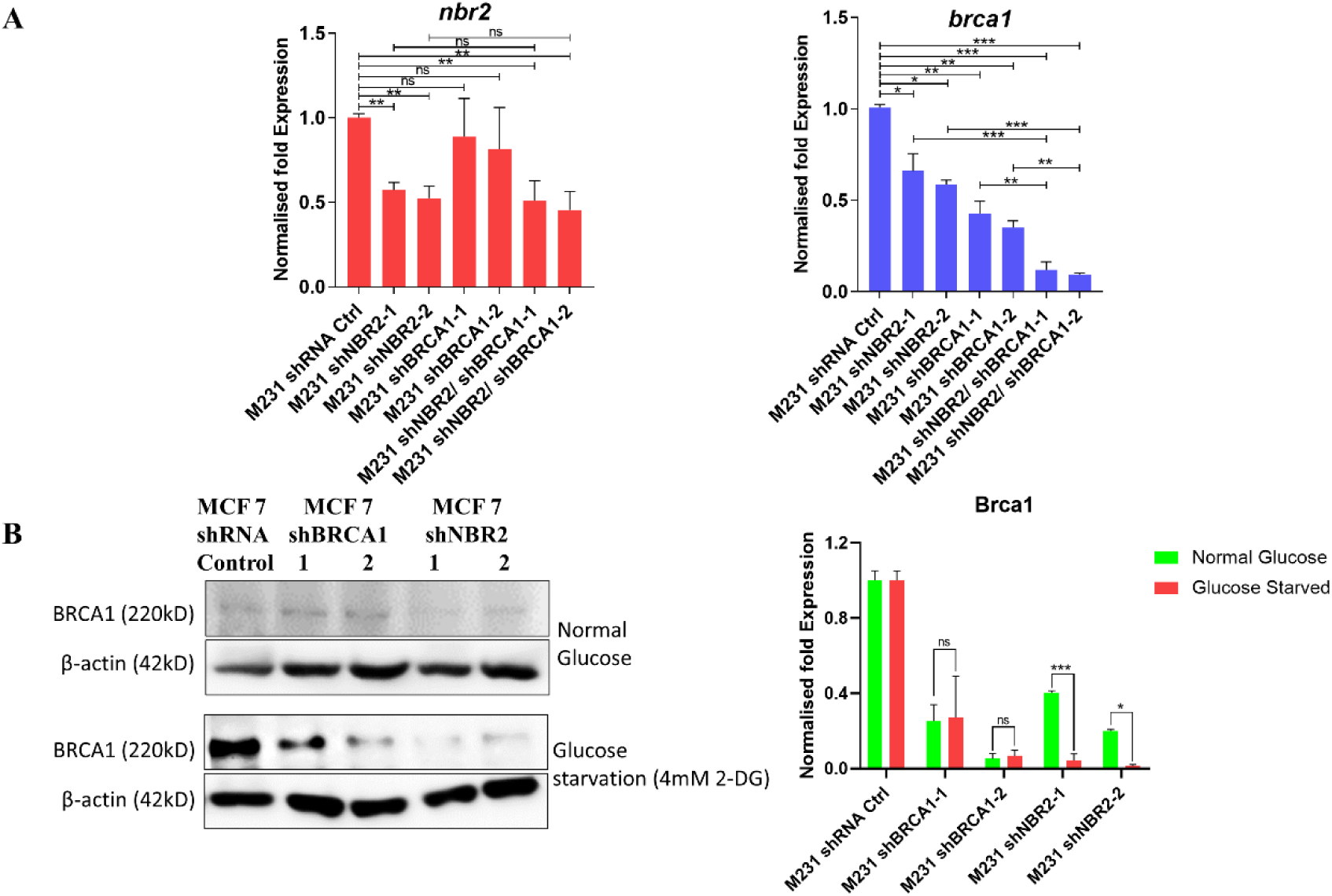
**(A)** NBR2 and BRCA1 knockdown studies by qRT-PCR show downregulation of BRCA1 mRNA upon NBR2 knockdown which is further enhanced in cells with double knockdown of NBR2 and BRCA1. **(B)** Immunoblot analysis show alleviated expression of BRCA1 in NBR2 knockdown condition which is further pronounced under glucose starvation condition.

### BRCA1 hypermethylation results in ectopic expression of β-hCG

Wildtype BRCA1 can bind to the β-hCG promoter [24] and regulate its expression. BRCA1 deficient cancers therefore can have high level of β-hCG expression which results in immune suppression, thereby inhibiting the anti-tumor response and leading to drug resistance[25, 28–30]. To examine whether the BRCA1 deficiency arising because of its promoter hypermethylation affects β-hCG, we analyzed β-hCG expression in cells with stably induced BRCA1 methylation. We observed significant overexpression of β-hCG in cells with both sgRNA 1 & 2 at RNA as well as protein level, which was approximately 3 times higher than BRCA1 knockdown cells at mRNA level (Fig 12). These results indicate that elevated level of β-hCG expression might be the reason for drug resistance observed in BRCA1 hypermethylated cancers [31, 32].

**Fig 12:**
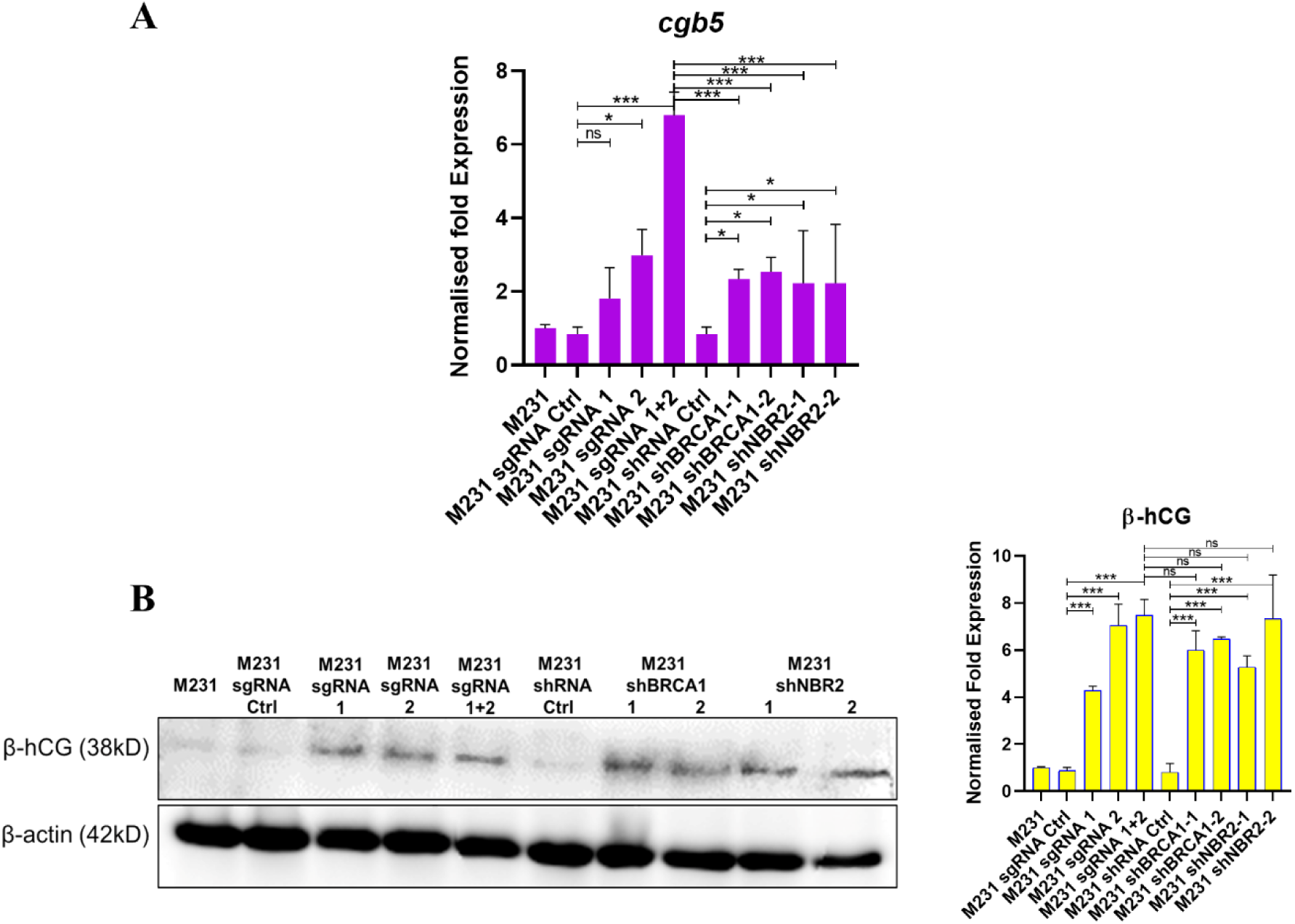
**(A)** β-hCG mRNA (CGB5) expression analysis shows upregulation of CGB5 levels in BRCA1 promoter α methylation induced M231 cells which is 87 folds higher than BRCA1 knockdown condition. **(B)** Immunoblot analysis shows ectopic expression of β-hCG in BRCA1 methylated cells which is significantly higher than BRCA1 knockdown condition.

### BRCA1 hypermethylation modulates hormone receptors allowing proliferation in the early stages and luminal to basal transformation at later stages of tumorigenesis

Considering the fact that a majority of BRCA1 hypermethylated as well as mutated cancers develop a TNBC phenotype, we investigated the effect of BRCA1 promoter methylation on the hormone receptors ER-α, PR and HER2. Our initial exploration into the breast cancer datasets from TCGA showed us a negative correlation between expression levels of ER, PR & HER2 and methylation levels of BRCA1 (Fig 13 I & II). Based on these results we analyzed the ER- α, ER-β and PR levels in the luminal cell line MCF 7. We observed alleviated levels of PR which was most significant in cells with both sgRNA1 & 2 (Fig 14A). However, to our surprise we found the ER-α expression to be upregulated in the methylation induced cells (Fig 14A). Meanwhile, ER-β which functions antagonistically to ER-α and has anti-proliferative effects was found to be downregulated (Fig 14A).

**Fig 13(I):**
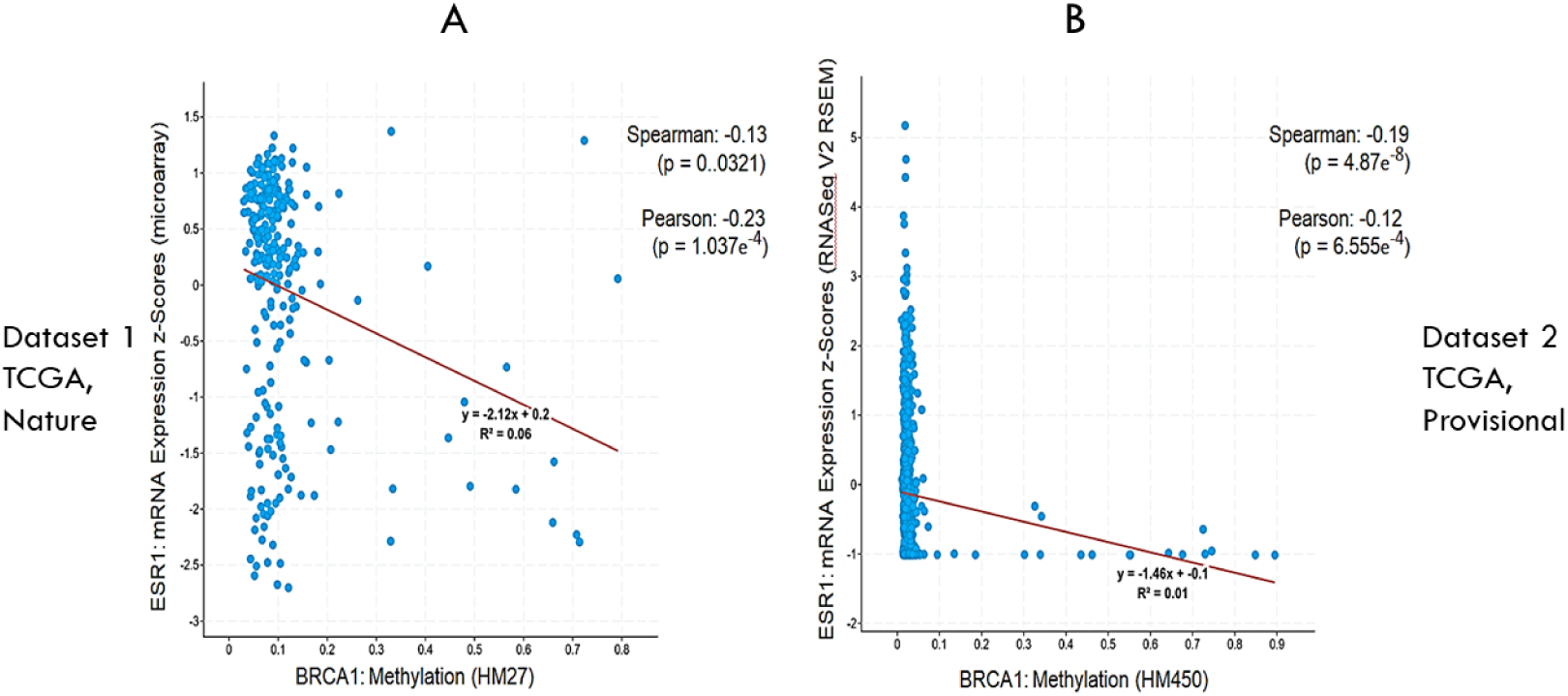
Plot A & B represent BRCA1 methylation VS ESR1 mRNA levels in the datasets Breast Invasive Carcinoma (TCGA, Nature 2012) and Breast Invasive Carcinoma (TCGA, Provisional) respectively.

**Fig 13(II):**
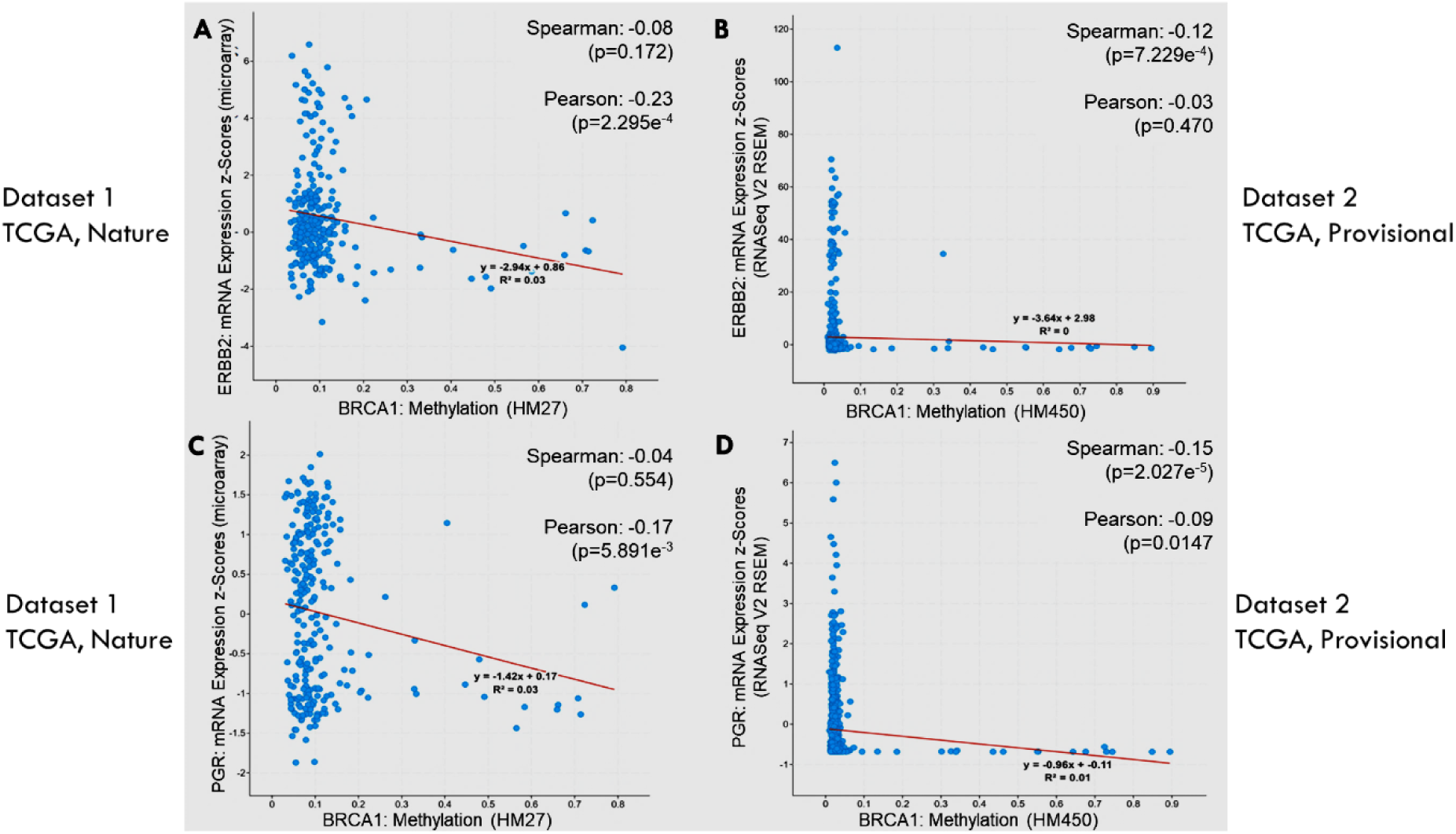
Plot A & B represent BRCA1 methylation VS ERBB2 mRNA levels in the datasets Breast Invasive Carcinoma (TCGA, Nature 2012) and Breast Invasive Carcinoma (TCGA, Provisional) respectively. Plot C & D represent BRCA1 methylation VS PGR mRNA levels in the same two datasets.

**Fig 14:**
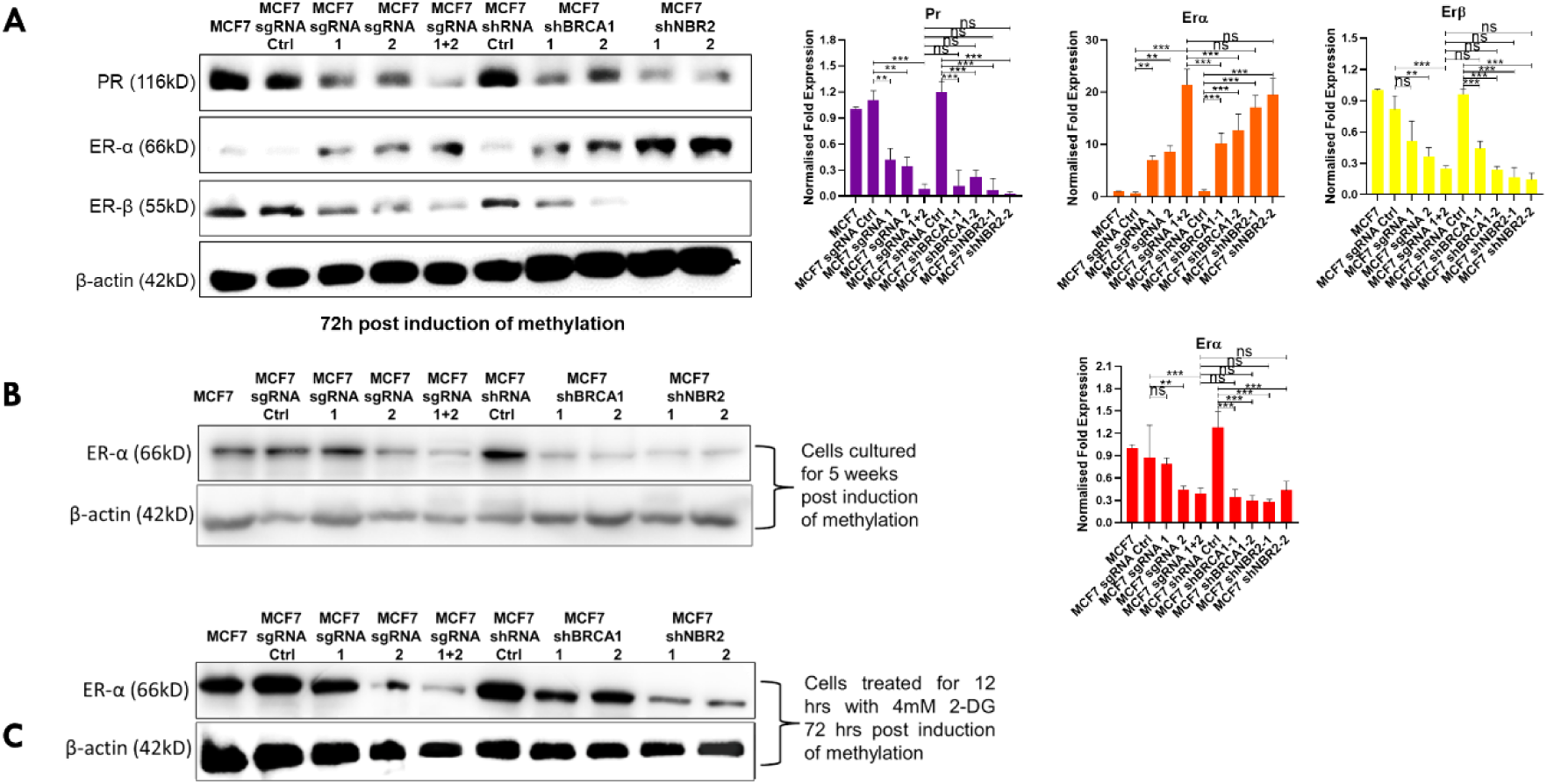
**(A)** Hormone receptor expression in MCF7 cells reveal downregulation of PR and an unexpected upregulation of ER-α and upregulation of ER-β levels in BRCA1 methylated cells. **(B)** Maintenance of methylation for 5 weeks post induction result in reduction in ER-α expression which was found to be higher in fig A. **(C)** Cells post 48 hrs of induction of methylation when subjected to glucose starvation mimic the late stage methylated cells and show alleviated ER- α expression similar to cells maintained for 5 weeks.

We hypothesized that this unusual expression pattern might be responsible for helping the cells survive and proliferate under the defunct DNA Damage Response in the absence of BRCA1 and this expression pattern may change at later stages of tumor formation. Following this idea, we cultured the cells for 5 weeks post induction of methylation and re-examined ER-α levels. As expected, we observed a downregulation of ER-α expression in both BRCA1 methylated and knockdown cells and NBR2 knockdown cells as well (Fig 14B). To mimic the later stages of tumor development, we also subjected the early stage methylated cells (48hrs post induction) to 12 hours glucose starvation and re-examined the ER-α expression. Concurrent with the previous findings, we observed a downregulation of ER-α levels in BRCA1 methylated, knockdown and NBR2 knockdown cells (Fig 14C). These results indicate that induction of BRCA1 promoter methylation leads to increased proliferation by modulating ER-α and ER-β expression initially, however results in the luminal to basal transformation at later stages of tumorigenesis.

### Reversal of Methylation leads to upregulation of BRCA1 and diminishes ER-α further

Unlike mutations, hypermethylation of BRCA1 is an epigenetic modification and it is reversible in nature. Therefore, we set out to investigate if reversal of induced methylation can reverse the tumorigenic effects it had on the cells over a period of time.

Since, our stable methylated and knockdown cells were under the control of a Tet-off promoter we treated the cells (with methylation induced and maintained for 5 weeks) with 1μg/ml Doxycycline to reverse the induction of methylation and examined the expression levels of BRCA1 and ER-α after 36hrs of Doxycycline treatment. We observed an increase in BRCA1 expression which was higher than the BRCA1 expression even in control cells (Fig 15). This overexpression of BRCA1 may be attributed to the fact that the induced cells were under a prolonged period of defunct DNA Damage Repair in absence of BRCA1 [33] which might have led to an accumulation of DNA damage and a temporary overexpression of BRCA1 as a response to DNA damage. But, to our amaze we found the ER-α levels to diminish even further upon revival of BRCA1 (Fig 15). We believe this unexpected downregulation is because of the transcriptional control that BRCA1 has over ER-α [34, 35] and we expect the ER-α levels to revive back to normal levels over time.

**Fig 15:**
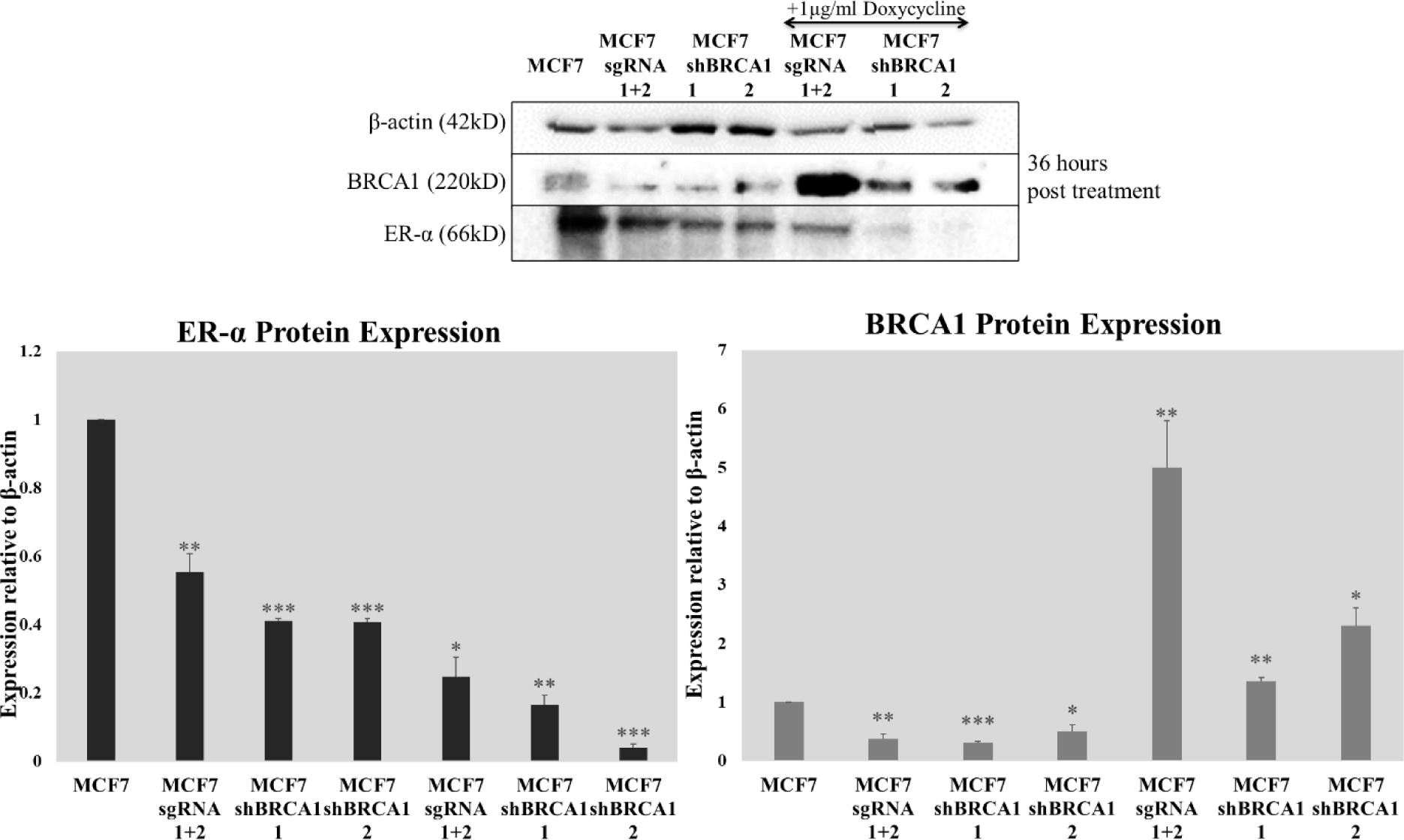
Reversal of BRCA1 methylation and knockdown (by 36 hours of doxycycline treatment) which was under the control of a tet-off promoter leads to overexpression of BRCA1 in response to DNA damage and causes the ER-α expression to drop further.

## DISCUSSION

This study makes an attempt to shed light on the reversible molecular play of an important epigenetic factor, promoter hypermethylation in driving the process of tumorigenesis. Although, BRCA1 promoter hypermethylation are more prevalent in sporadic breast and ovarian neoplasms, they do have a role to play in inherited cancer cases as well. Evidence shows that epimutations like promoter hypermethylation can be passed on through generations and can lead to cancer predisposition [36–38]. Recent studies have also shown the presence of BRCA1 hypermethylation in pregnant adults and its maternal transfer to their newborns [39, 40]. Another recent report suggests that constitutive BRCA1 hypermethylation may be responsible for early onset breast cancers [41]. All these impute a higher emphasis to its role in tumor predisposition and progression.

In this study we have seen that induction of hypermethylation in the BRCA1 promoter α in MDA-MB-231 cells lead to an abnormal shift in the balance between the two alternate BRCA1 transcripts. This is also accompanied by a lower BRCA1 expression which is probably because of the switching of transcription from α to β promoter, since the α promoter remains unavailable for the transcription factors to bind. The concurrent downregulation of the tumor suppressing lncRNA, NBR2 is an augmenting factor that works as a confederate with BRCA1 hypermethylation in tumor progression as NBR2 downregulation not only imparts invasive property to cells [18, 42, 43], but also generates a feedback loop which increments the silencing of BRCA1. Although NBR2 downregulation is associated with aggressive forms of breast tumors, certain class of drugs show better prognosis under NBR2 deficiency [44]. This suggests that biguanides like metformin (widely used for diabetes treatment) and phenformin can be employed to target NBR2 deficient BRCA1 methylated tumors and achieve a better anti-tumor response. Interestingly, a recent study shows suppression of colorectal cancer progression by upregulation of NBR2 upon Curcumin treatment [45] which is already known to be a potent hypomethylation agent, causing reversal of hypermethylation in breast [46], gastric [47] and colorectal cancer [48]. This indicates that curcumin may have the ability to suppress breast cancer development and improve prognosis by reversal of BRCA1 hypermethylation and revival of NBR2 expression. However, another recent report suggests that BRCA1 hypermethylated breast tumors gain resistance to DNA damaging drugs by reversal of promoter hypermethylation [31], which makes the future of hypomethylating cancer therapies uncertain and open for further studies.

Overexpression of β-hCG is a key driver of aggressive manifestations in BRCA1 mutated cancers which is accompanied by several levels of immune modulations and hypermethylation of BRCA1 promoter developed a kindred outcome. The similarity in phenotypes of BRCA1 hypermethylated cancers with BRCA1 mutated cancers which is termed as BRCAness [49], may also be a consequence of such common genes and pathways affected in the tumorigenic process. Our findings also suggest that development of β-hCG inhibitors may serve to be effective in combinatorial therapies to alleviate the aggressiveness and drug resistance in both BRCA1 mutated and hypermethylated tumors.

As BRCA1 has a transcriptional hold over β-hCG, the expression of β-hCG in BRCA1 hypermethylated cells was an expected outcome. However, this expression of β-hCG being several folds higher in BRCA1 hypermethylated cells than BRCA1 knockdown cells (at mRNA level) is very striking and is an indicative of involvement of other transcriptional regulators of β-hCG. One of the key hormones that is known to control β-hCG expression in placenta is Progesterone. Reports have shown that progesterone can inhibit hCG synthesis by inhibiting the accumulation of α and β-hCG mRNA [50]. This evidence suggests that under reduced expression of PR the progesterone hormone activity might be affected, thereby leading to accumulation of β-hCG mRNA in BRCA1 methylated cells. The negative status of PR is reported to be more frequent in malignant tumors [51] and is often reported to be associated with high proliferative indices as well [52]. Studies have also shown an association between progesterone receptor negative status and tumor relapse[53–55]. Thus, the reduced level of PR in combination with high expression of β-hCG can be considered as one of the indispensable reasons as well as markers for tumor aggressiveness in the BRCA1 methylated cells.

The selective and differential modulation of ERs in different stages of tumor development demonstrate how cancer cells use the same molecules for their growth at one stage and shed them at a later stage to escape the targeted anti-tumor chemotherapies. ER-α negative tumors that express ER- β show better prognosis to endocrine therapies like adjuvant tamoxifen treatment [56] and lack of ER- β is associated with resistance to tamoxifen therapy [57] & early relapse[58] in ER-α negative cancers. The low levels of ER-β in methylated cells suggests that tamoxifen or ER-β antagonist mediated chemotherapy may not be very effective in targeting these tumors. Clinically, like BRCA1 mutation screening and risk evaluation, screening of BRCA1 hypermethylation, in combination with NBR2 & β-hCG levels as markers could prove to be very beneficial as well as expedient in early diagnosis of breast cancer, evaluation of risk of future breast neoplasms and development of targeted treatment strategies in sporadic and even inherited breast cancers.

## Notes

### Competing Interest Statement

The authors have declared no competing interest.

### Summary of Updates

Data for shRNAs have been shown for two different shRNA constructs for better replicability. shRNA mediated NBR2 knockdown data has been used wherever applicable for a more clear understanding of the data

